# GlnK facilitates the dynamic regulation of bacterial nitrogen assimilation

**DOI:** 10.1101/127662

**Authors:** Adam Gosztolai, Jörg Schumacher, Volker Behrends, Jacob G Bundy, Franziska Heydenreich, Mark H Bennett, Martin Buck, Mauricio Barahona

## Abstract

Ammonium assimilation in *E. coli* is regulated by two paralogous proteins (GlnB and GlnK), which orchestrate interactions with regulators of gene expression, transport proteins and metabolic pathways. Yet how they conjointly modulate the activity of glutamine synthetase (GS), the key enzyme for nitrogen assimilation, is poorly understood. We combine experiments and theory to study the dynamic roles of GlnB and GlnK during nitrogen starvation and upshift. We measure time-resolved *in vivo* concentrations of metabolites, total and post-translationally modified proteins, and develop a concise biochemical model of GlnB and GlnK that incorporates competition for active and allosteric sites, as well as functional sequestration of GlnK. The model predicts the responses of GS, GlnB and GlnK under time-varying external ammonium level in the wild type and two genetic knock-outs. Our results show that GlnK is tightly regulated under nitrogen-rich conditions, yet it is expressed during ammonium run-out and starvation. This suggests a role for GlnK as a buffer of nitrogen shock after starvation, and provides a further functional link between nitrogen and carbon metabolisms.

## Introduction

To adapt to the highly variable nutrient conditions in their natural habitat, microorganisms have evolved a complex intracellular circuitry coupling signal transduction, membrane transport, gene expression and metabolism. A widely conserved example is the ammonium assimilatory system in *Escherichia coli*, which coordinates the uptake of nitrogen with other pathways involved in carbon assimilation and maintenance of cellular energy status (1–3). External ammonium *(E. coli*’s preferred nitrogen source) is assimilated into glutamate and glutamine in two steps: first, ammonium and intracellular glutamate (GLU) are converted into glutamine (GLN) in a reaction catalysed by the enzyme glutamine synthetase (GS); second, glutamine is combined with *α*-ketogluterate (*α*-KG), a product of carbon metabolism, to yield two molecules of glutamate (for a net production of one GLU molecule) in a reaction catalysed by the enzyme glutamate synthase (GOGAT) (Fig. 1).

**Figure 1:**
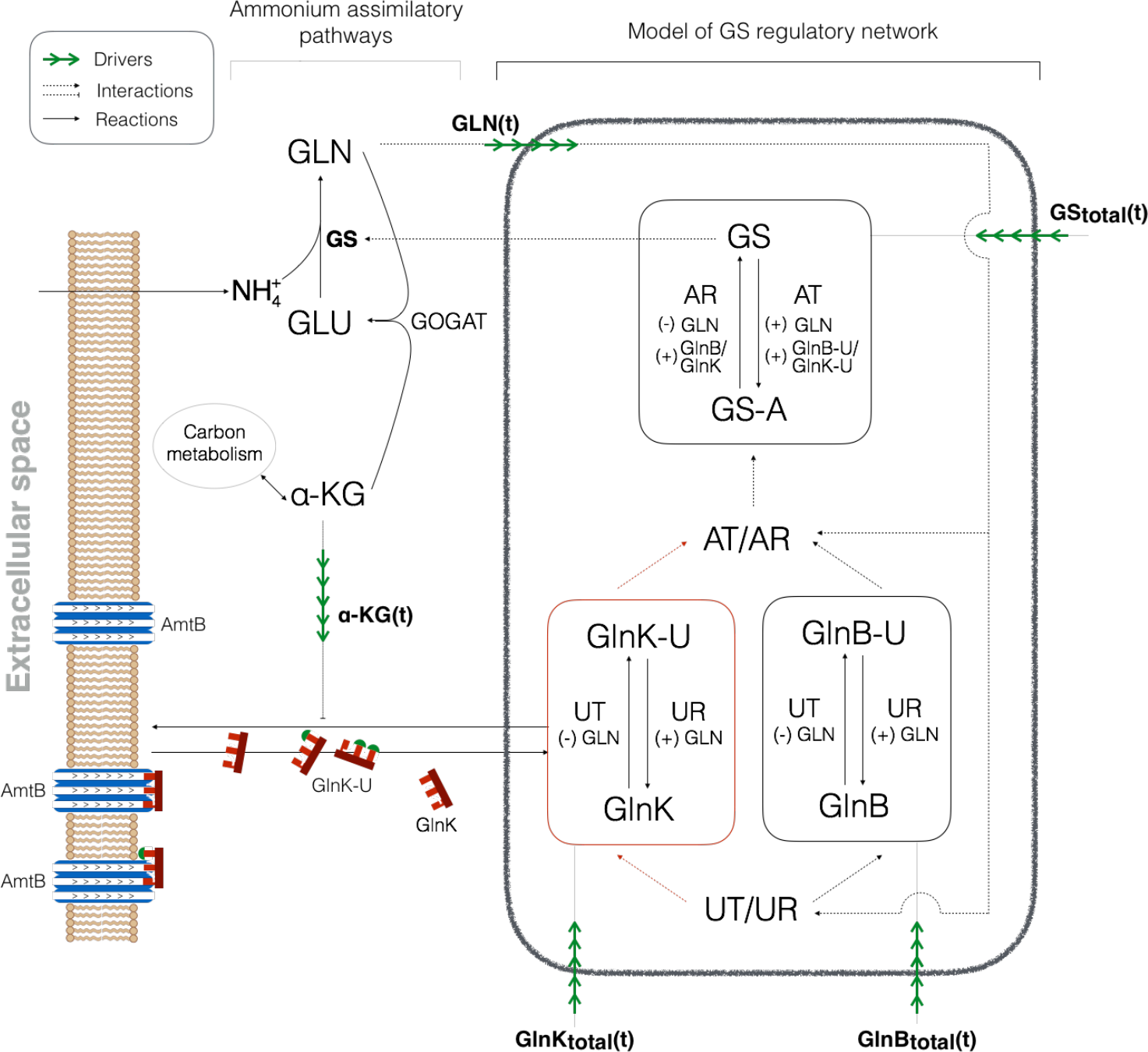
Summary of the ammonium assimilatory system in *E. coli*. Under nitrogen limited conditions, intracellular ammonium is assimilated almost exclusively through an enzymatic reaction catalysed by GS. The activity of GS is controlled by the enzymes in the black box, which are the object of our model, as a function of the carbon (*α*-KG) and nitrogen (GLN) states of the cell. Our model describes the post-translational states of GlnB, GlnK and GS, which are modified by the enzymes AT/AR and UT/UR. The total concentrations of GlnB, GlnK and GS, which reflect the transcriptional and translational responses of the cell, are measured experimentally and taken as drivers for the purposes of the model. Glutamine is sensed by UT/UR and transmitted via two complementary branches of signal transduction regulated by GlnB and GlnK. The focus of this study is GlnK (red box). Trimers of GlnK form a complex with the membrane-bound transport protein AmtB, with the effect of blocking active ammonium import. During run-out and starvation, high levels of *α*-KG inhibit AmtB-GlnK complex formation, thus enabling active transport. Importantly, AmtB-GlnK complex formation sequesters GlnK, changing the amount accessible by UT/UR and AT/AR. The nitrogen and carbon information contained in the total and PTM levels of GlnB and GlnK are integrated with the level of GLN by the enzyme AT/AR to regulate the activity of GS.

Although glutamine and glutamate are both key intermediates towards the production of amino acids and nucleotides, their wider roles in cellular metabolism are different. Whereas glutamate is the nitrogen source for most (88%) reactions in the cell (1), glutamine constitutes the primary signal controlling the activity of enzymes in the ammonium assimilation system. As a result, their regulatory requirements are different: homeostatic regulation of glutamate is essential to maintain growth rate, whereas glutamine levels must adjust rapidly to reflect cellular nitrogen status while at the same time remaining within physiologically acceptable bounds. In particular, when nutrient-rich conditions are suddenly restored after a period of starvation, the cell must be able to avoid glutamine shock (4, 5).

The modulation of GS activity is the primary means by which *E. coli* achieves these regulatory targets in the face of changing nitrogen levels. On one hand, as extracellular ammonium is used up during growth, the activity and abundance of GS is raised so as to maintain the growth rate as long as possible. On the other hand, when favourable conditions are suddenly restored, GS can be ‘turned off’. This nonlinear response of GS is controlled by the bifunctional enzyme adenylyl-transferase/adenylyl-removase (AT/AR), which can reversibly adenylylate (inactivate) or deadenylylate (activate) independently each of the 12 monomers that conform GS, thus modulating GS activity in a graded manner (6). The activity of AT/AR is in turn regulated allosterically by glutamine and by two highly similar signal transduction proteins, GlnB and GlnK (encoded, respectively, by the *glnB* and *glnK* genes) (2, 7). Importantly, GlnB and GlnK share a source of upstream regulation, themselves being reversibly uridylylated by a second bifunctional enzyme, uridilyl-transferase/uridylyl-removase (UT/UR), which enables GlnB and GlnK to control the enzyme AT/AR (8). Specifically, under glutamine limitation GlnB and GlnK are mostly uridylylated, leading to increased AR activity and inhibition of AT activity (Fig. 1). In addition to regulating AT/AR, GlnB and GlnK also regulate the expression of GS and bind *α*-KG, communicating with carbon metabolism and signal transduction pathways (2).

Due to the apparent redundancy of having two signalling proteins (GlnB and GlnK) relaying the glutamine state of the cell, previous models (9–12) have ignored GlnK regulation of AT/AR, it being less potent than GlnB. This simplification is supported by mutational studies of the receptor interaction domains of GlnB and GlnK (13) and by *in vitro* studies (14). From this viewpoint, the role of GlnK is mostly circumscribed to regulating the membrane-bound channel protein AmtB, whose gene *(amtB*) lies in the same operon as *glnK*, causing *glnK* and *amtB* expression to be directly correlated (15–17).

However, various experiments suggest that GlnK also acts as an effective regulator of AT/AR. For instance, GlnB deficient strains (i.e., Δ*glnB* mutants) exhibit unimpaired growth (18) with regulated GS (de)adenylylation (19), suggesting that GlnK can substitute for GlnB. It has also been found that GlnK does interact with AT/AR *in vitro* (14), albeit relatively weakly, and binds *α*-KG, ATP and ADP similarly to GlnB (20, 21). Importantly, *glnK* expression is induced during nitrogen limitation, whereas *glnB* is expressed constitutively (19). Taken together, these observations raise questions about the *in vivo* regulation of AT/AR and about a complementary role for GlnK in nitrogen assimilation to provide not only added redundancy but also a flexible and rapid response to the consequences of variations in external ammonium levels.

To establish the relevance of GlnK in nitrogen regulation during external nitrogen variations, we combined experiment with modelling. We measured the growth rate (Fig. 2) and the *in vivo* temporal response of relevant metabolites, proteins and post-translational modification (PTM) protein states during ammonium run-out, starvation and a subsequent ammonium shock (Fig. 3). Measurements of the wild type (WT) strain were complemented with experiments with two genetic knock-outs: Δ*glnB* (no GlnB) and Δ*glnK* (no GlnK), as shown in Fig. 5.

**Figure 2:**
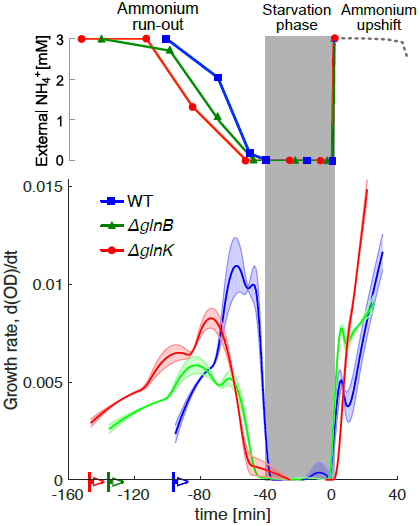
Experimental time series of growth rate of the WT and the two mutants (Δ*glnB* and Δ*glnK*) during ammonium run-out followed by a step change of ammonium. During the starvation phase (marked by the grey shaded area), the growth is halted. The time series of the three strains were aligned such that the onset of starvation coincides. The three strains were introduced to the medium at different times, as indicated by the coloured arrows. The lines and symbols represent averages and the coloured bands represent ±s.e (n=3). The growth rate was computed using a Savitzky-Golay filter applied to the experimental measurements of the optical density (OD) over the course of the experiment.

**Figure 3:**
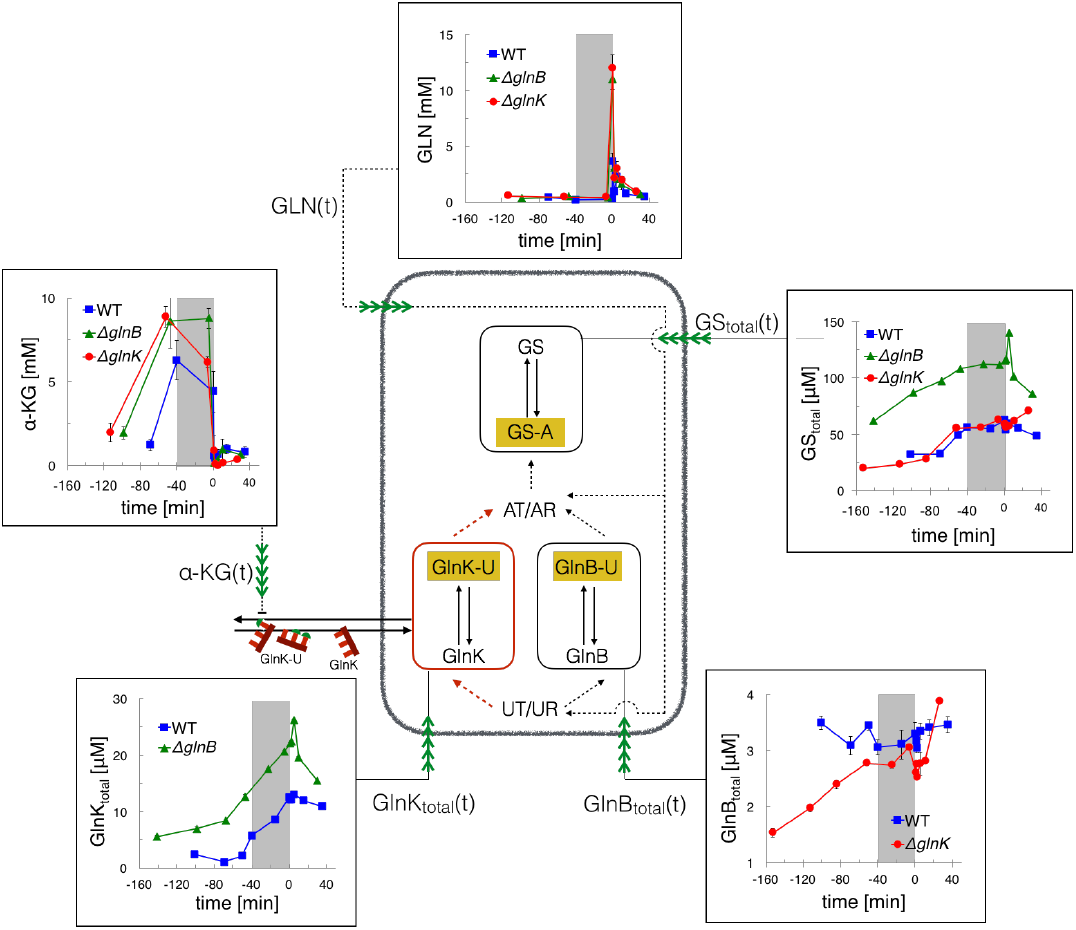
Experimental time series of total monomeric protein (GlnB, GlnK and GS) and metabolite (glutamine and *α*-KG) concentrations for the WT (blue), Δ*glnB* (green) and Δ*glnK* (red) strains. Symbols and error bars represent average protein or metabolite concentrations ±s.e (n=3). The grey shaded area marks the starvation period. The lines are guides to the eye. This data was used to drive the model (pictured) of the PTM of the three proteins (highlighted in orange). Time-course measurements of the modified proteins GS-A, GlnB-U and GlnK-U were also taken experimentally and are presented in Fig. 5.

**Figure 4:**
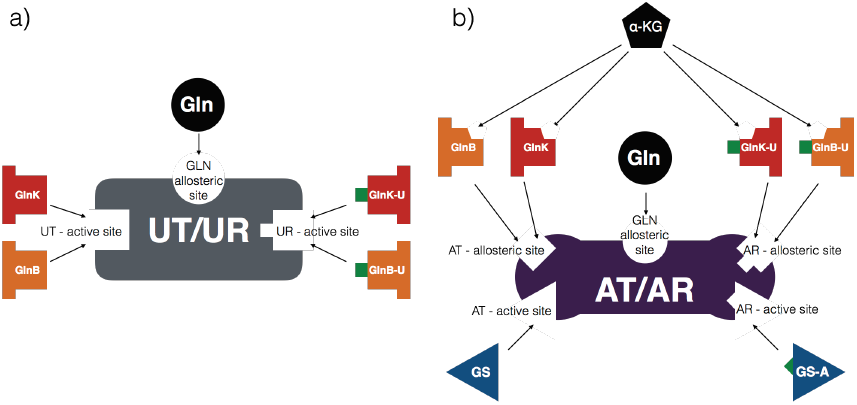
Schematic illustration of the structure of UT/UR and AT/AR. a) UT/UR has two distinct active sites: one binds GlnB-U and GlnK-U, whereas the other binds the unmodified GlnB and GlnK species. Hence GlnB and GlnK are competing substrates for UT/UR. In addition, UT/UR is allosterically regulated by glutamine, which activates the UR and inhibits the UT activity. b) AT/AR has two separate active sites: one binds GS, the other GS-A. It also has three distinct allosteric sites: the first binds GlnB-U and GlnK-U and induces AR excitatory and AT inhibitory responses; the second binds GlnB and GlnK and induces AT excitatory and AR inhibitory responses; the third binds GLN and modulates the rate of AT and AR responses. In addition, *α*-KG can bind to both the modified and unmodified GlnB/GlnK proteins, affecting their interaction with AT/AR.

**Figure 5:**
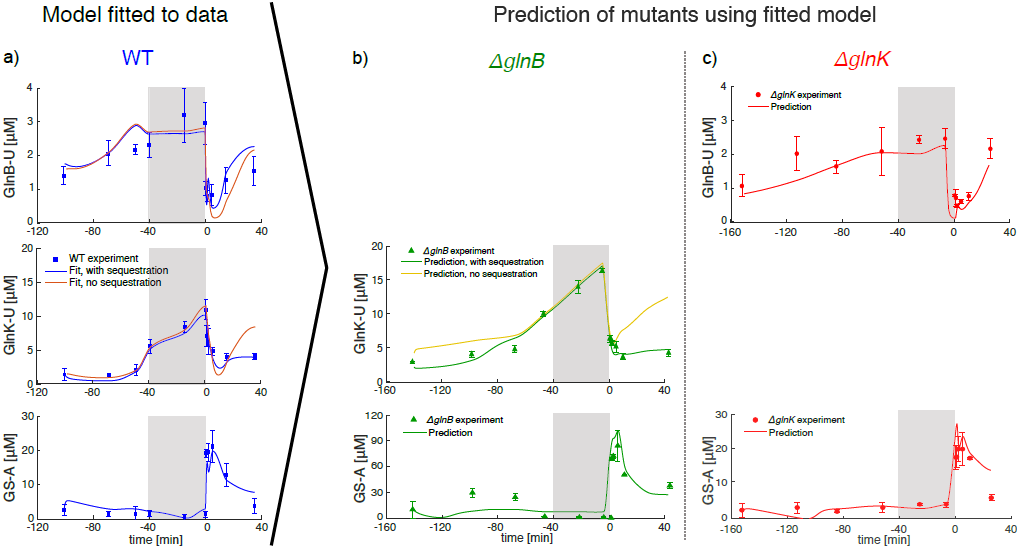
Time responses of PTM protein levels in the WT and two knock-out mutants. a) Time responses of GlnB uridylylation, GlnK uridylylation and GS adenylylation in the WT strain (experimental measurements and fitted model, with and without sequestration). b-c) Experimental measurements of GlnB uridylylation, GlnK uridylylation and GS adenylylation for knock-outs: b) Δ*glnB* (green) and c) Δ*glnK* (red), compared in both cases to the dynamics predicted by our model fitted only to the WT data. The symbols and error bars represent measured average protein concentrations ±s.e (n=3). The solid lines correspond to fits and prediction of our model, as indicated. The grey shaded area marks the starvation period. The model with sequestration provides a significantly better fit according to information criteria for model selection (see text).

Our experiments show that growth rate and ammonium uptake are not impaired by the removal of either GlnB or GlnK (Fig. 2) suggesting that GlnB and GlnK conform two complementary branches of nitrogen signal transduction regulating AT/AR. We also find that GlnK plays a limited role in regulating GS expression and adenylylation during ammonium run-out and starvation. However, GlnK levels rise steeply during run-out, exceeding those of GlnB four-fold during starvation (Fig. 3). This suggests a functional role of GlnK, post-starvation, until normal ammonium conditions are restored.

To confirm this hypothesis, we developed a concise, mechanistic model of GS regulation (Fig. 1). We show that competition between GlnB and GlnK for UT/UR active sites and AT/AR allosteric sites, as well as functional sequestration of GlnK are necessary to explain the *in vivo* uridylylation and adenylylation dynamics in the WT and to predict those in the two mutants. Our model predicts the dynamic response of GS adenylylation levels in the mutants, indicating that GlnK can substitute for GlnB. Moreover, we find that GlnK is a potent regulator of AT/AR post-starvation, when GlnK levels are high relative to GlnB. Our results agree with the role of GlnK as a regulator of the ammonium transporter AmtB upon nitrogen upshift (15, 16), and with the decreased viability of *glnK* mutants following extended ammonium starvation (22). Hence our work suggests a role of GlnK as an anticipatory buffer molecule to help avoid glutamine shock: GlnK is expressed before and during starvation in order to temporarily alleviate excess ammonium import and assimilation when ammonium becomes available again.

## Materials and methods

### Strains and growth conditions

Experiments were performed with *E. coli* K12 (NCM3722) (23). NCM3722Δ*glnB* in-frame deletions were obtained through P1 phage transduction (24), using strain JW2537 with the *glnK* knock-out from the Keio collection as the donor strain (25). NCM3722Δ*glnB* was verified by genomic locus sequencing. Pre-cultures were grown in Gutnick minimal media, supplemented with 10 mM NH_4_Cl, 0.4% (w/v) glucose and Ho-Le trace elements. Main cultures were inoculated in same media but with limiting 3 mM NH_4_Cl, resulting in NH_4_Cl starvation at mid log phase. Ammonium depletion and OD600 were determined at defined time points during the course of experiments as described by (26). Following one generation time (40 min) of growth arrest, NH_4_Cl was added to obtain a concentration of 3 mM (upshift).

### Metabolite and protein measurements

Glutamine and *α*-ketoglutarate were quantified using liquid chromatography-mass spectrometry (LC-MS) and nuclear magnetic resonance (NMR), as described previously (26, 27). Total abundance and uridylylation level of GlnK were determined using multiple reaction monitoring mass-spectrometry (MRM-MS) (28) with absolute standard protein quantification (PSAQ) (26, 29). Briefly, isotopically labelled GlnK protein standard was purified following IPTG induced over-expression in *E. coli* harbouring JW0440 from the ASKA(-) collection (*glnK*) (30), grown in the presence of labelled L-Arginine (^13^C_6_, ^15^N_4_) and L-Lysine (^13^C_6_, ^15^N_2_). Ni-affinity purified GlnK standard purity was judged visually by SYPRO Ruby Protein Gel Stain (Invitrogen) stained SDS-PAGE to be >90% and GlnK standard concentration determined. Isotopic labelling efficiency was determined by MRM-MS to be 100%. Total GlnK abundances were determined by the ratio of MRM-MS signals of GlnK unlabelled/labelled signature peptides, excluding the uridylylation site peptide (Table S1 in the Supporting Material) and intracellular concentrations calculated (26). GlnK uridylylation levels were directly measured by the abundance of the uridylylated GlnK peptide GAEYS(**UMP**)VNFLPK compared with the non-uridylylated peptide GAEYSVNFLPK. The GlnK protein concentrations derived from non-uridylylated peptides correspond well with GlnK protein concentrations derived independently from the sum of uridylylated and non-uridylylated peptide GAEYSVNFLPK (Fig. S8 in the Supporting Material). A similar procedure is used for quantifying GlnB, GS and their PTMs (27).

### Description of the model

The schematic of the system regulating nitrogen assimilation in *E. coli* is shown in Fig. 1. We use three coupled non-linear ODEs to describe the dynamics of the monomeric concentrations of uridylylated GlnB, uridylylated GlnK and adenylylated GS:

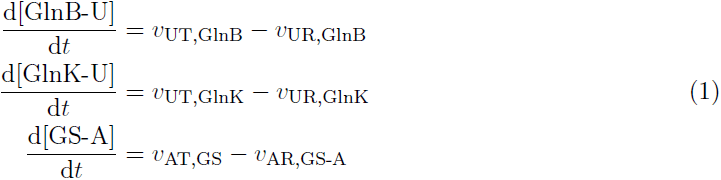
where the right hand side contains the balance of fluxes corresponding to (de)uridylylation and (de)adenylylation of the corresponding proteins. These fluxes contain several kinetic parameters and depend non-linearly on the concentrations of substrates, products and allosteric effectors. We now describe these terms in detail.

#### Equations for UT/UR

The UT and UR activities in the bifunctional enzyme UT/UR are treated as independent, unidirectional reactions obeying Michaelis-Menten kinetics. Due to the specificity of UT/UR active sites (31), GlnB and GlnK are modelled as competing substrates. Our model considers only the experimentally measured monomeric concentration of the uridylylated species, and disregard the multimeric nature of GlnB and GlnK, since (de)uridylylation of GlnB monomers is non-cooperative (32). Glutamine (GLN) acts as an allosteric activator to UR and an inhibitor to UT activities (8). These can be encapsulated in the following rate equations (see Section S1 in the Supporting Material for derivations):

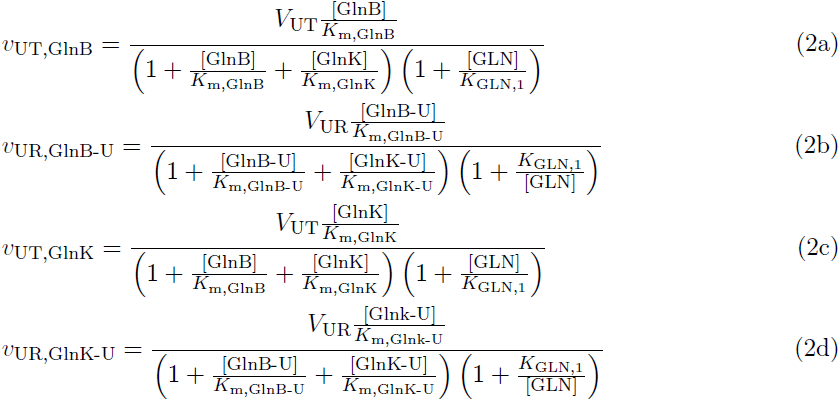

Here *V*_*_ and *K*_m,_* are the maximal rates and Michaelis constants of the (de)uridylylation reactions, and *K*_GLN,1_ is the binding affinity of GLN to the allosteric site of UT/UR. The concentration of GlnB (and similarly GlnK) is obtained from the conservation relation:

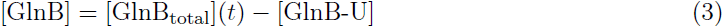
where the total concentration is a time-varying experimental measurement (Fig. 3).

#### Sequestration of GlnK

We assume that GlnK-AmtB complex formation regulates the amount of GlnK accessible to other reactions, and is controlled by *α*-KG following a Hill-type form (17):

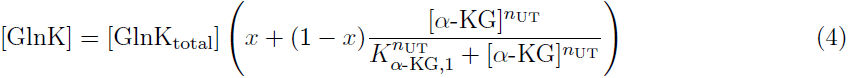
where [GlnK_total_] and [GlnK] are the total and accessible concentrations of GlnK, respectively; *K*_*α*-KG,1_ is the dissociation constant for *α*-KG; *n*_UT_ is the Hill-coefficient; and *x* is the minimum concentration of accessible GlnK, which indicates the ratio of AmtB to GlnK. If [*α*-KG] ≪ *K*_*α*-KG,1_, then [GlnK] = *x*[GlnK_total_], and the maximal amount of GlnK is sequestered; if [*α*-KG] ≫ *K*_*α*-KG,1_, then [GlnK] ≈ *x*[GlnK_total_], so no GlnK is sequestered.

#### Equations for AT/AR

We split the activity of AT/AR into independent AT and AR reactions, both obeying Hill-type kinetics. We consider that GlnB-U/GlnK-U and GlnB/GlnK are pairwise competing for allosteric sites, while glutamine can bind independently to a third allosteric site (7, 21, 33). The binding of GlnB and *α*-KG are synergistic, and that of GlnB-U and *α*-KG are antagonistic (34) leading to the following activation parameters:

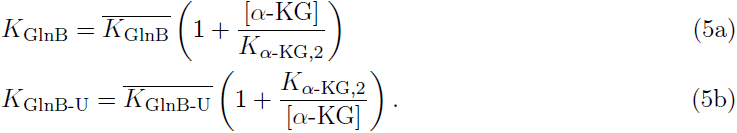

Similar expressions are used for GlnK and GlnK-U. As above, we only describe GS-A monomers (without explicit GS-A complexes) (9, 35, 36). We neglect retroactivity from GlnB and GlnK on *α*-KG, since the latter is two orders of magnitude more abundant, and from AT/AR on GlnB and GlnK, since GlnK level is expected to be significantly higher (7). These assumptions result in the following rate equations (see Section S2 in Supporting Material for a detailed derivation):

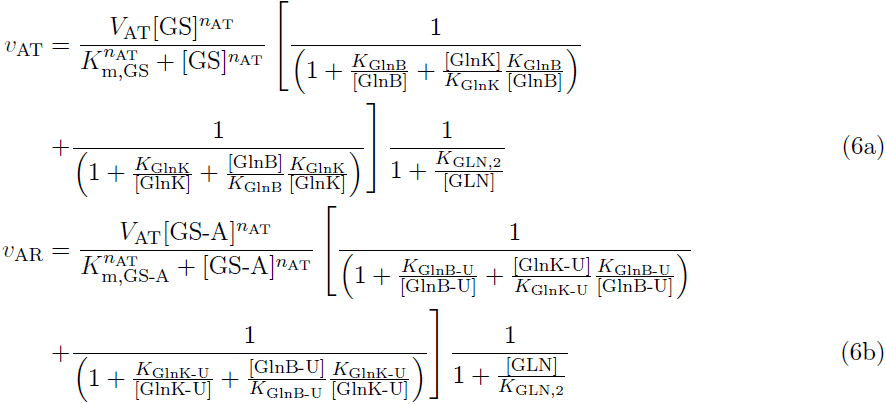
where *V*_*_ and *K*_m,*_ are the maximal enzyme rate and Michaelis constants of (de)adenylylation, *K*_*_ are the binding affinities of GlnB(-U), GlnK(-U) and GLN to the allosteric sites of AT/AR and *n*_AT_ is the Hill-coefficient. A summary of these interactions is shown in Figure 4.

Simulations were performed in Matlab. An SBML format of the model is included in the Supporting Material where we refer the reader for further details.

### Model parameters: fitting and sensitivity analysis

The parameters were taken from the literature whenever possible (6 out of 21) and the rest (15 out of 21) were fitted to the 36 time points of the WT data (Table 1).

**Table 1:**
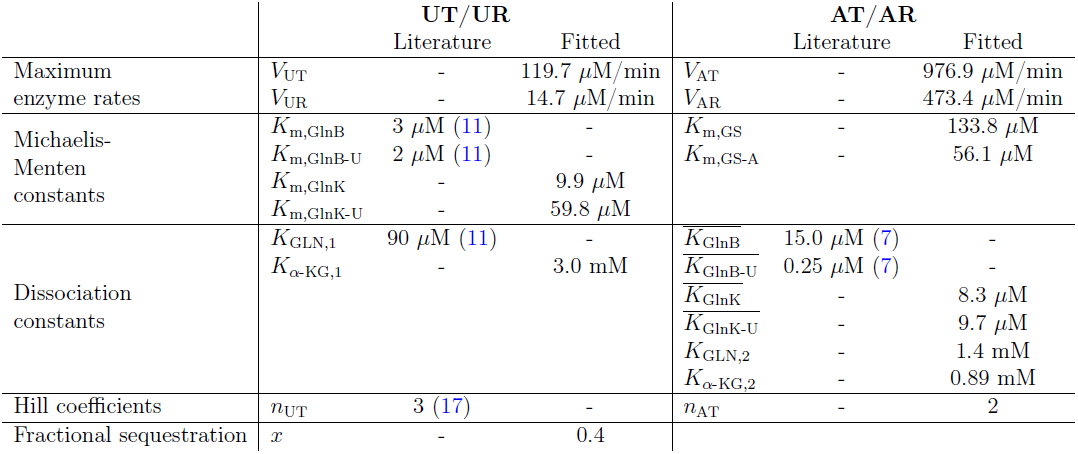
Model parameters

|  | UT/UR |  |  | AT/AR |  |  |
| --- | --- | --- | --- | --- | --- | --- |
|  |  | Literature | Fitted |  | Literature | Fitted |
| Maximum enzyme rates | $V_{UT}$ | - | 119.7 $\mu\text{M}/\text{min}$ | $V_{AT}$ | - | 976.9 $\mu\text{M}/\text{min}$ |
| | $V_{UR}$ | - | 14.7 $\mu\text{M}/\text{min}$ | $V_{AR}$ | - | 473.4 $\mu\text{M}/\text{min}$ |
| Michaelis-Menten constants | $K_{m,\text{GlnB}}$ | 3 $\mu\text{M}$ (11) | - | $K_{m,\text{GS}}$ | - | 133.8 $\mu\text{M}$ |
| | $K_{m,\text{GlnB-U}}$ | 2 $\mu\text{M}$ (11) | - | $K_{m,\text{GS-A}}$ | - | 56.1 $\mu\text{M}$ |
| | $K_{m,\text{GlnK}}$ | - | 9.9 $\mu\text{M}$ | | | |
| | $K_{m,\text{GlnK-U}}$ | - | 59.8 $\mu\text{M}$ | | | |
| Dissociation constants | $K_{\text{GLN},1}$ | 90 $\mu\text{M}$ (11) | - | $\overline{K_{\text{GlnB}}}$ | 15.0 $\mu\text{M}$ (7) | - |
| | $K_{\alpha\text{-KG},1}$ | - | 3.0 mM | $\overline{K_{\text{GlnB-U}}}$ | 0.25 $\mu\text{M}$ (7) | - |
| | | | | $\overline{K_{\text{GlnK}}}$ | - | 8.3 $\mu\text{M}$ |
| | | | | $\overline{K_{\text{GlnK-U}}}$ | - | 9.7 $\mu\text{M}$ |
| | | | | $K_{\text{GLN},2}$ | - | 1.4 mM |
| | | | | $K_{\alpha\text{-KG},2}$ | - | 0.89 mM |
| Hill coefficients | $n_{UT}$ | 3 (17) | - | $n_{AT}$ | - | 2 |
| Fractional sequestration | $x$ | - | 0.4 | | | |

To assess the robustness of the model and to evaluate the relevance of the parameters to be fitted, we performed a global sensitivity analysis using the eFAST algorithm (37). This technique considers uncertainty in combinations of parameters and quantifies the deviation they introduce over the whole time series. See Section S3 in the the Supporting Material for details of the method and its results.

To fit the parameters of the model to the measured time series, we used the Squeeze-and-Breathe evolutionary Monte Carlo optimisation algorithm (38). This algorithm minimises a cost function *J*, which is the sum-of-squares deviation of the fractional PTM levels from the measurements, weighted by the relative errors. Mathematically,

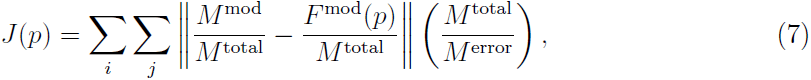
where *p* denotes the vector of parameters; *M* and *F* denote measured and fitted, respectively; and the superscripts *mod, total* and *error* denote total, modified and error, respectively. The parameter values are presented in Table 1, rounded to the nearest decimal found in the confidence interval given by the distribution of optimised parameters obtained with the Squeeze-and-Breathe method (see Section S4 in the Supporting Material for details).

## Results

### *In vivo* measurements of *E. coli* temporal responses to nitrogen starvation and upshift

We measured concentrations of relevant metabolites (*α*-KG and GLN) and active/inactive PTM states of the proteins GlnB, GlnK and GS in response to dynamic variations in external ammonium (see Materials and Methods). We use this novel experimental approach to measure PTM states (26, 27) in order to link metabolic status and regulatory mechanisms.

Three *E. coli* cultures (a WT strain and two isogenic strains with deleted *glnK* or *glnB*) were inoculated into minimal media with a (limiting) concentration of 3 mM ammonium (see Materials and Methods). The growing cultures depleted the external nitrogen levels during the run-out period eventually leading to the starvation period measured from the point when growth stopped and extending for 40 minutes (Fig. 2). The starvation period was followed by a sudden 3 mM ammonium upshift to investigate the shock response (0 min to 40 min). We sampled the cultures at specific external ammonium concentrations during run-out and at specific time points during starvation and upshift, and measured the metabolite and protein concentrations. The corresponding time courses are plotted on Fig. 3.

### *In vivo* regulation of GS by GlnK

First, we investigated whether our experiments provide evidence for GlnK controlling the level of GS expression. We found no discernible difference in total GS level between Δ*glnK* and WT (Fig. 3), indicating that GlnK is redundant to regulate GS expression in the WT. In addition, we found that GlnK and GS were two-fold over-expressed in Δ*glnB* (Fig. 3). This observation provides an *in vivo* quantification of the reduced regulatory effect of GlnK on GS expression, which has been previously evidenced in genetic studies (14).

Next, we asked whether GlnK plays a relevant role as a regulator of GS adenylylation. The response of the Δ*glnB* strain to external ammonium variations was consistent with GlnK substituting for GlnB in its role to control AT/AR (18). Although the growth of Δ*glnB* was slower than the WT during ammonium run-out, it was comparable to Δ*glnK* (Fig. 2), indicating that, relatively speaking, ammonium uptake was not limiting. We further found that GS was abruptly adenylylated upon nitrogen upshift in all strains (Figs. 5c,e,g), including the Δ*glnB* mutant, despite higher GS levels in the latter (Fig. 3). This suggests that, after starvation, GlnK is a potent activator of GS adenylylation. In contrast, GS was not fully deadenylylated in Δ*glnB* (Fig. 5e and Fig. S6a in the Supporting Material) under ammonium rich conditions (unlike in WT and in Δ*glnK*, see Fig. 5c,e), showing that GlnK might be less efficient in activating the AR activity. Indeed, GlnK was almost fully uridylylated (>95%) during starvation (see Fig. S6c in the Supporting Material), consistent with the dominant GlnK species under starvation being the fully uridylylated GlnK trimer, GlnK_3_-U_3_, which is known to be a poor activator of deadenylylation *in vitro* (39).

Unlike in Δ*glnB*, we observed no evidence of GlnK regulated GS adenylylation in the WT during ammonium run-out and starvation, the GS-A levels being similar between the WT and Δ*glnK* (Fig. 5c,g). Despite this purported redundancy, the expression and uridylylation of GlnK are already induced when ammonium levels begin to decrease (19) (Fig. 3 and 5b). Since GlnB expression is constitutive, we find that during starvation the concentration of GlnK (12.5 *μ*M) largely exceeds that of GlnB (3.3 *μ*M) (Fig. 3) and similarly GlnK-U (11 *μ*M) largely exceeds GlnB-U (2.9 *μ*M) (Fig. 5a,b). Thus intracellular glutamine sensing by UT/UR operates before ammonium concentration becomes suboptimal for growth.

Taken together, these findings suggest a role for GlnK to prepare for a buffered return to subsequent normal nitrogen conditions after starvation, by regulating the inactivation of GS. Our measurements show that the Δ*glnK* strain grows at a higher rate after ammonium upshift (Fig. 2). We hypothesise that this difference is due to additional ammonium being imported via the membrane-bound ammonium channel AmtB, which accumulates during nitrogen starvation and whose activity becomes unimpeded at low levels of GlnK.

To understand further the interaction between GlnB and GlnK during nitrogen runout, starvation and upshift, we constructed a mathematical model motivated by our *in vivo* dynamic measurements of protein (total and PTM) and metabolite concentrations. To date, the use of directly determined PTM data in models of regulation of GlnB and GlnK has been lacking, yet such modification directly determines their functionality. We now present an inclusive modelling approach to link GlnB and GlnK activity via PTMs to metabolic status and control of nitrogen assimilation processes.

### Mathematical modelling of the enzymatic network of nitrogen signalling and regulation

In view of the experimental data, we built an ordinary differential equation (ODE) model to describe the dynamics of PTM states, i.e., the (de)uridylylation of GlnB and GlnK and the (de)adenylylation of GS in response to temporal inputs (drivers). An ODE model was adopted due to the large copy numbers observed—the smallest being GlnB with an average of ~ 10^3^ molecules per cell. We based our equations on the enzyme mechanisms reported in the literature with the aim of a parsimonious model, only including terms with an observable effect in our experiments. The resulting model remains mechanistic with a moderate number of parameters, facilitating direct comparison to experiments and parameter fitting.

The terms of the ODE model are derived from the biochemical details of competition for the active and allosteric sites, which is inherent to the uridylylation and adenylylation reactions, and from the formation of complexes between GlnK and AmtB. In addition, we use five measured input drives to isolate the responses pertaining to transcription and other parts of the *E. coli* metabolism. These include concentrations of metabolites GLN and *α*-KG, which act as enzyme effectors and reflect the metabolic state of the cell, and the total concentrations of GlnB, GlnK and GS, which reflect the relevant transcriptional responses of the cell (Fig. 3). We further assume that key cofactors (e.g., ATP, NADPH) remain at homeostasis under our experimental conditions (10), so that their concentrations can be approximated to be constant and can thus be absorbed into the kinetic parameters of the equations.

The model parameters include: binding affinities (*K*_d_) of various effectors to active and allosteric sites of the enzymes; Michaelis constants (*K*_m_) and maximum enzyme rates (*V*_max_); and parameters representing AmtB-GlnK complex formation (Table 1). Whenever possible we used parameters from the biochemical literature, and we fitted the rest to the WT experimental time-courses using the Squeeze-and-Breathe algorithm (38). To confirm that the parameters capture meaningfully the experimental observations, we also performed a global sensitivity analysis over the time series (37). For details, see Materials and Methods and the Supporting Material.

To make predictions for the Δ*glnB* and Δ*glnK* mutants, we set the corresponding reaction rates to zero, fixing all other parameters at values fitted to the WT time-course data. We then generate predicted time series for the PTM protein states under the experimentally measured drives. These predictions are compared to the independently obtained measurements.

### Modelling the mechanism of GlnB and GlnK uridylylation

The bifunctional enzyme UT/UR has two independent active sites catalysing uridylylation (UT) and deuridylylation (UR), which are highly specific for GlnB and GlnB-U, respectively (31); it also has an allosteric site for glutamine (Fig. 4a). Due to the similarity between GlnB and GlnK, we assume similar specificity of UT/UR active sites to GlnK/GlnK-U, yet with different binding affinities. Hence we model GlnB and GlnK uridylylation as two monocycles, GlnB-UT/UR and GlnK-UT/UR, which operate in parallel but not independently, since the total amount of enzyme is distributed between them at any given time. We further consider that the binding of glutamine inhibits the uridylylation of GlnB/GlnK and activates the deuridylylation of GlnB-U/GlnK-U (8) (Eq. 2, Materials and Methods).

As shown in Fig. 5a, our model captures accurately the excitatory and inhibitory effects in the measured GlnB uridylylation under different conditions in the WT without the need for modelling ternary complexes explicitly as in the more complex model in Ref. (12) (Fig. S5 in the Supporting Material). The parameters fitted to the WT data show that the affinities of GlnK and GlnK-U to AT/AR (*K*_m_ = 9.9*μ*M and 59.8*μ*M, respectively) are significantly lower than those of GlnB and GlnB-U (*K*_m_ = 3*μ*M and 2*μ*M, respectively), which are known experimentally (11). Hence GlnK only affects nitrogen assimilation when its concentration is several times higher than GlnB, in agreement with previous *in vitro* findings (14).

Although this model allows us to fit well the overall WT time measurements, it cannot capture the GlnK-U level in response to the ammonium upshift after starvation (Fig. 5a). Moreover, the model over-predicts the level of GlnK-U in the Δ*glnB* knock-out (Fig. 5b). In the next section we discuss an additional mechanism, which is necessary to make accurate predictions of the observed GlnK-U levels.

### Sequestration of GlnK is necessary to predict uridylylation levels

To accurately describe GlnK uridylylation dynamics, we take into account the interaction of GlnK with AmtB. The membrane-bound channel protein AmtB facilitates ammonium active import to the cell. AmtB channels are blocked when AmtB forms a complex with trimers of deuridylylated GlnK above 20-50 *μ*M external ammonium concentrations (16). Hence AmtB-facilitated transport is negligible in our experiments before complete run-out. However, the formation of complexes between AmtB and GlnK does affect the dynamical responses, since it provides a mechanism for the *sequestration* of GlnK and GlnK-U at the membrane (40). Note that while fully uridylylated trimers cannot bind AmtB (15), our measurements show that the uridylylation level of GlnK is around 30% after starvation (see Fig. S6c in the Supporting Material) suggesting that the GlnK trimers will be mostly mono-uridylylated, i.e. one uridylylated GlnK per GlnK trimer, on average. In our model, we assume that both fully deuridylylated GlnK trimers and partially uridylylated GlnK trimers may be functionally sequestered to the membrane. However, unlike the fully deuridylyated GlnK trimers, partially uridylylated GlnK trimers do not block AmtB channel activity.

We model AmtB-GlnK complex formation by allowing a proportion of the total GlnK to be sequestered with a Hill-function dependence (Eq. 4, Materials and Methods) on the concentration of *α*-KG (16). Note that, since high levels of *α*-KG level promote the dissociation of the AmtB-GlnK complex, sequestration only occurs during external ammonium abundance, i.e., when the *α*-KG levels are low (Fig. 3).

The model with sequestration fitted to the WT data is able to capture the GlnK-U levels after nitrogen upshift (Fig. 5a). To justify the necessity of the two additional parameters (*K*_*α*-KG,1_, *x*) involved in GlnK sequestration, we use the fitted model to predict the uridylylation states of GlnK and GlnB in the Δ*glnK* and Δ*glnB* knock-outs, and compare them to the measured time-courses (Fig. 5b,c). This approach has been used before to develop systems biology models (41, 42). Our predictions agree with the measurements: without sequestration, the model overpredicts the levels of GlnK-U before starvation (time before −40 min) and after nitrogen upshift (time after 0 min) in the Δ*glnB* mutant (Fig. 5d). To quantify this, we compute the error of both models (with and without sequestration) against the data (7), and rank both models according to two information theoretic criteria (AICc, BIC), which penalise the additional parameter complexity against any improvement in the error of the model. The extended model with sequestration is clearly selected by both information criteria (see Table S2 in the Supporting Material) lending further basis for the need for a GlnK sequestration term during nitrogen excess.

The higher growth rate observed in the Δ*glnK* strain after starvation (Fig. 2) is also consistent with GlnK sequestration. Since *amtB* is still present in this strain, AmtB is expected to accumulate during nitrogen starvation, which provides unrestricted ammonium import and higher growth rate in the short term.

### GlnK can regulate AT/AR in the absence of GlnB

Our results so far indicate that GlnK not only competes with GlnB as a substrate or UT/UR, but also provides a link between active membrane transport and carbon metabolism through the interaction with *α*-KG. To further test this mechanism, we asked: is GlnK sufficient on its own to regulate the adenylylation enzyme AT/AR in the absence of GlnB? If so, one would expect that the predicted GlnK-U levels in the Δ*glnB* strain should account correctly for GS-A levels and, hence, for the activity of GS in nitrogen assimilation.

To test this hypothesis, we extended our model to include GS adenylylation. Similarly to UT/UR, the two antagonistic activities of AT/AR are modelled as independent unidirectional reactions, justified by structural studies showing the two active sites to be well separated (33), making interaction between the AT and AR domains less likely (Fig. 4b).

We model different facets of AT/AR regulation as found in the literature (Fig. 4b). Firstly, AT and AR activities are inhibited by the products, GS-A and GS, respectively (7). Secondly, in the absence of allosteric effectors AT/AR has no observable activity (7). From this state, the rates of AT and AR actitivites are regulated by five allosteric effectors, which no not affect the binding rates of substrates (21). In particular, GLN, GlnB and GlnB-U may bind to three distinct sites (7, 21, 33). As above, we assume that GlnK(-U) has the same specificity as GlnB(-U) but different affinity, so that GlnB and GlnK compete for one allosteric site, and GlnB-U and GlnK-U compete for the other. Finally, to account for intramolecular signalling (33), we allow the activation parameters of GlnB, GlnB-U, GlnK and GlnK-U to depend on *α*-KG (Eqs. 5, Materials and Methods). We encapsulate these terms into the reaction schemes (Eq. 6), by generalising the classical allosteric activation model (43) for two competing effectors (see Section S2 in the Supporting Material for details).

The additional equation for GS-A was fitted to the WT data and captures well the observed variability (Fig. 5a). To avoid overfitting, we keep fixed all parameters related to UT/UR and sequestration, and fix the insensitive activation parameters 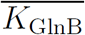 and 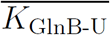 to the values reported by (7) (Table 1, Materials and Methods and Section S3 in the Supporting Material). In contrast, the activation parameters 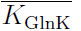 and 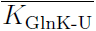 are relatively insensitive in the WT, yet essential to predict GS-A levels in the Δ*glnB* strain. The fitted activation constant *K*_GLN,2_ for AT/AR is in the range reported in Ref. (7) and the Hill coefficient *n*_AT_ is found to be as reported in Ref. (35). Note also that the activation constants of GlnK and GlnK-U for AT/AR are much higher than those of GlnB and GlnB-U, consistent with their observed lower affinity (14).

To assess the model fitted to the WT data, we predict the level of GS-A for the two knockouts in response to the measured drives. The predicted responses agree with the measured GS-A levels in both strains (Fig. 5b,c). The GlnB-deficient strain (Δ*glnB*) is able to regulate GS activity in response to external changes in ammonium. The prediction for the Δ*glnB* strain underestimates the GS-A level before starvation—we consider the possible source for this discrepancy in the Discussion. Given that GS regulation depends on the fine balance between GlnB/GlnB-U and GlnK/GlnK-U together with the level of GLN and *α*-KG, our model demonstrates that GlnK can regulate GS under varying ammonium conditions.

## Discussion

We have investigated the role of GlnK in the dynamic regulation of ammonium assimilation from a systems perspective, and have argued that paradoxical observations about GlnK stem from the subtle interplay of regulatory mechanisms on two levels: transcriptional and post-translational. To decouple both levels of regulation, we collected concurrent *in vivo* time-courses of metabolite, total protein and PTM protein concentrations in response to time-varying external ammonium levels. We analysed the data in conjunction with a mechanistic model derived from biochemical principles. The combined analysis allowed us to fit the WT dynamics of PTMs, and to used the fitted model to predict the dynamic responses of experimental data from genetic knock-outs, shedding light on experimental ambiguities about the regulatory role of GlnK.

We found that competition between GlnB and GlnK for the bifunctional enzyme UT/UR together with GlnK sequestration by AmtB describe the uridylylation responses in the WT and can predict the response of two mutants under dynamic ammonium perturbations. AmtB-GlnK complex formation not only blocks ammonium import (16), but also rapidly sequesters GlnK, whereas the less abundant GlnK limits ammonium assimilation by facilitating more rapid deuridylylation of GlnK-U (Fig. 5) and thus higher GS-A levels. Thus GlnK dampens the assimilatory response to ammonium upshift, perhaps to avoid excess ammonium uptake causing the depletion of the glutamate pool (4, 5). The fact that sequestration is regulated by *α*-KG (a product of the carbon metabolism), suggests that nitrogen shock responses could also help to avoid untoward impacts on carbon status.

Our experiments showed that during ammonium rich conditions, GlnK is present at low level and does not influence GS production and adenylylation, which is instead regulated by the more potent GlnB. However, due to the induction of *glnK* during ammonium run-out, GlnK outnumbers GlnB four-fold during starvation (Fig. 3). To examine the capability of GlnK for AT/AR regulation after ammonium upshift, we developed a mathematical model of GS adenylylation based on allosteric competition between GlnB and GlnK for AT/AR. The model predicts accurately the abrupt increase in GS-A levels after ammonium upshift in both the Δ*glnB* and Δ*glnK* strains (Fig. 5) showing that both GlnB and GlnK are independently sufficient for effective AT/AR regulation. Our model slightly underestimates the GS-A levels before starvation in the Δ*glnB* strain. Since the model was fitted to the WT, where both GlnB and GlnK are present, this mismatch may indicate unaccounted-for interactions which are disrupted when deleting the *glnB* gene, e.g., GlnB and GlnK forming heterotrimers (39, 44, 45).

To check the relevance of GlnK regulation of AT/AR, we used our model to simulate two ‘in silico’ cellular scenarios. Firstly, we considered our model for a ‘computational strain’ where GlnK does not interact with AT/AR (by setting 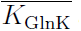 ≫ [GlnK] and 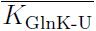 ≫ [GlnK-U], such that the terms involving GlnK and GlnK-U drop out in Eq. 6), and hence AT/AR is regulated only by GlnB. In this scenario where the regulatory role of GlnK is removed, the predicted GS-A levels in response to ammonium upshift after starvation are reduced (Fig. 6a), thus leading to an increase in GS activity and larger ammonium assimilation. This is consistent with the proposed role of GlnK as a buffer to ammonium shock, through the adenylylation (inactivation) of GS. Noting that the level of GS-A does not change during run-out and starvation, we also used our model to predict the response of another ‘computational strain’ (Fig. 6) where the expression of *glnK* is constitutive and unregulated (by fixing [GlnK_total_] as a constant). In this scenario, if GlnK abundance is kept fixed at the late-starved level (26 *μ*M), GS-A levels remain elevated during run-out and starvation. This simulation supports the notion that GlnK must be kept down-regulated during nitrogen abundance, since early *glnK* expression would lead to reduced ammonium assimilation due to increased GS-A levels. During ammonium run-out and starvation, *amtB* (and conjointly *glnK*, being on the same operon) is expressed so as to increase ammonium uptake via AmtB channels. The resulting higher levels of GlnK induced by starvation serve also to control potential overshoots in ammonium levels following starvation not only by blocking AmtB channels, but also by directly reducing GS activity through adenylylation.

**Figure 6:**
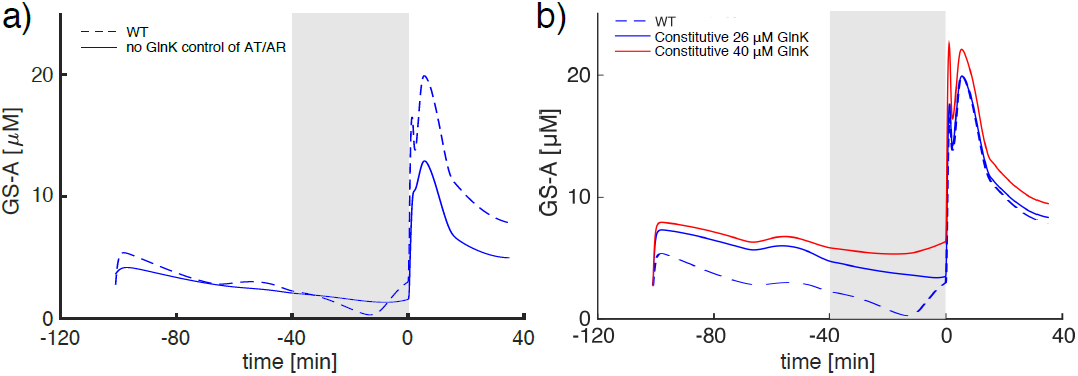
Predictions of the model for the GS-A levels of two ‘computational strains’. a) In the first ‘strain’, AT/AR is regulated only by GlnB but not by GlnK, yet GlnK still regulates AmtB and competes with GlnB for UT/UR. Before starvation, the levels of GS-A are not affected by GlnK due to low GlnK abundance. If GlnK is assumed not to regulate AT/AR, the level of GS-A after ammonium upshift is reduced, implying increased ammonium assimilation. Hence the high level of GlnK induced during starvation in the WT acts as a buffer to prevent overshoots in ammonium assimilation by regulating GS activity. b) In the second ‘strain’, the expression of *glnK* (and conjointly of *amtB*) is constitutive (unregulated). In order to avoid the overshoot due to ammonium upshift, a high level of GlnK is necessary before starvation. This over-abundance of GlnK pre-starvation leads to higher GS-A levels, and hence suboptimal ammonium uptake. In each figure, the grey shaded area indicates the ammonium starvation period; the dashed line shows the original model for the WT data (same as in Fig. 5); the solid lines are predictions of the model for the respective ‘computational strains’.

Beyond nitrogen regulation, our work exemplifies the need for *in vivo* experimental time series data in developing predictive models. Time-course inputs together with classic genetic knock-outs provide tools to gain insight into the dynamic aspects of regulation in signalling pathways. Although large-scale biochemical models can be powerful, their applicability is many times hampered due to the large number of unknown parameters they contain, which need to be fitted to scarce data. Here we derived a small set of mathematical modules commensurate with the observed dynamics, and justified their relevance by a global parameter sensitivity analysis. We used a recently developed parameter fitting algorithm (38), which is especially appropriate for time-courses, to fit the WT data, and we then used the fitted model to predict out-of-sample dynamical observations from the genetic knock-outs.

Although we have focussed here on *E. coli*, with the dual GlnB/GlnK proteins, we expect nitrogen assimilation and transport to be more strongly coupled in bacteria containing only one GlnB-like protein (e.g. cyanobacteria, azotobacter), enabling the tight coordination of carbon and nitrogen metabolisms. Similarly, patterns of sequestration may be a biophysical regulatory mechanism operating widely in other contexts, and understanding its spatiotemporal dynamics in the cell would help link gene and metabolic regulation responses to biophysical feedback to environmental inputs (46).

## Author contributions

AG, MBa, MBu and JS designed the study. AG performed the analysis under the supervision of MBa and MBu. JS designed and coordinated experiments performed by VB, JGB, FH, and MBe. AG, MBa, JS and MBu wrote the manuscript, with inputs from other authors.

## Acknowledgements

We thank Mariano Beguerisse-Diaz for his intellectual contributions about parameter fitting. This work was funded by a BBSRC LoLa grant (BB/G020434/1). AG acknowledges funding through a PhD studentship under the BBSRC DTP at Imperial College London (BB/M011178/1). M. Barahona acknowledges funding from the EPSRC (EP/N014529/1 and EP/I032223/1).

## Conflict of interest

The authors declare that they have no conflict of interest.

